# Public Cohort Analysis Identifies Thyroglobulin Variants as Hypothyroidism Risk Factors

**DOI:** 10.1101/2025.11.21.689591

**Authors:** Jake N Hermanson, Andrew D Hudson, Lars Plate

## Abstract

Hypothyroidism is a prevalent endocrine disorder characterized by insufficient thyroid hormone (T3T4) production. Thyroglobulin (Tg) serves as the prohormone for T3 and T4 production, with many variants of uncertain clinical significance due to genetic diversity in the Tg gene. We leveraged the large-scale *All of Us* biobank to investigate the disease association of prevalent yet undercharacterized Tg variants. We related variant presence to thyroid-stimulating hormone levels and levothyroxine (LT4) usage as proxies for thyroid function. This identified R152H, Q870H, A993T, P1012L, and P1494L variants linked to increased LT4 usage and decreased thyroid function, while the R320C variant was associated with decreased thyroid function. Molecular characterization in Fisher rat thyroid cells revealed decreased secretion efficiency of R152H, Q870H, and R320C variants. Affinity purification-mass spectrometry demonstrated that secretion-deficient variants showed higher engagement with the protein homeostasis network, indicating protein quality control defects as the pathophysiology mechanism. In contrast, secretion-competent A993T and P1494L variants showed elevated interactions with degradation and antigen-presentation pathways, suggesting an alternative pathophysiology possibly linked to Hashimoto’s disease, an autoimmune condition with overproduction of autoantibodies that target thyroid proteins. In support, participants carrying the A993T or P1494L variants had elevated anti-TPO antibody levels. We estimate ∼150,000 US individuals currently taking levothyroxine could benefit from precision medicine targeting these variants, with ∼100,000 carrying Q870H. Our findings highlight the power of combining large public biobank data with molecular characterization to understand Tg genotype-to-phenotype relationships. Q870H represents a candidate for molecular therapies to restore secretion, offering precision medicine beyond LT4 replacement therapy.

**Significance Statement:** Hypothyroidism affects millions of Americans who typically receive levothyroxine hormone replacement therapy, yet some patients continue experiencing symptoms despite treatment. Correlating genomic and health data from the large-cohort *All of Us* biobank identified specific variants in the thyroglobulin gene, which produces thyroid hormones, that contribute to thyroid dysfunction through distinct biological pathways. Molecular characterization and interactomics revealed that some variants impair hormone secretion due to protein misfolding, while others associate with immune presentation that may link to autoimmune thyroid disease. The Q870H variant emerged as a promising target for precision medicine, potentially benefiting ∼100,000 Americans currently taking levothyroxine. This research demonstrates how combining large-scale genomic data with molecular characterization can identify new therapeutic targets beyond standard hormone replacement therapy.

## Introduction

Hypothyroidism is a prevalent disease characterized by the inability to biosynthesize enough triiodothyronine (T3) and thyroxine (T4), which are important thyroidal hormones for many metabolic and developmental processes.^1–4^ Production of these hormones is regulated by the hypothalamus-pituitary thyroid (HPT) axis. First, thyrotropin-releasing hormone (TRH) is secreted from the hypothalamus to induce the secretion of thyroid-stimulating hormone (TSH) from the pituitary gland.^5^ From there, TSH acts on thyroid cells to activate genes responsible for the biosynthesis of T3T4. After sufficient accumulation of T3T4, a negative feedback loop inhibits TRH and TSH secretion.^5^ The current standard of care for hypothyroidism patients is treatment with levothyroxine (LT4), a chemically synthesized form of T4. It is estimated that approximately 7% of the US population currently takes LT4, making it one of the top 10 prescribed drugs.^6,7^ Even though hypothyroidism is treatable with LT4, improper dosing of LT4 can lead to potential side effects that mimic hyperthyroidism, such as increased heart rate, anxiety, and difficulty sleeping.^8^ Patients taking LT4 exhibit an imperfect balance in T3 to T4 ratio, as dosing with LT4 only provides T4, while T3 has to be generated peripherally from T4.^9^ Since T3 is the more bioactive hormone, this imbalance may explain why some patients still report symptoms of hypothyroidism even with treatment of LT4.^4^ Additionally, circulating levels of T3T4 are variable depending on the time of day, season and illness, among other variables.^5,10^ A precision medicine strategy that allows the HPT axis to self-regulate T3T4 levels would circumvent these LT4 treatment complications.

Hypothyroidism disease etiology is heterogeneous, caused by genetic, autoimmune, and environmental conditions.^11^ Introduction of iodine into salt has greatly reduced the occurrence of environmentally caused hypothyroidism cases.^11^ Most congenital hypothyroidism cases occur from abnormal thyroid development (dysgenesis) or defective hormone synthesis (dyshormonorgenesis).^12^ Sequence variants in the biosynthesis genes are frequently the cause of dyshormonogenesis. Another disorder disrupting T3 and T4 biosynthesis is Hashimoto’s disease – an autoimmune condition where antibodies against Tg and/or TPO are produced, targeting the thyroid and disrupting its function.^13^

The biosynthesis of T3 and T4 in the thyroid is coordinated by many proteins and enzymes, such as thyroglobulin (Tg) and thyroid peroxidase (TPO), among other genes.^14^ Tg serves as the prohormone to T3T4 production. It is a 330 kDa multidomain protein that is secreted into the follicular lumen of thyroid glands. Here, it is first iodinated by TPO. Subsequently, an auto-catalyzed coupling can occur between a diiodotyrosine (DIT) and monoiodotyrosine or two DIT residues to generate the T3 or T4 precursors, respectively.^4,15^ Many Tg variants that result in hypothyroidism are misfolding variants retained in the endoplasmic reticulum (ER).^16–23^ Without proper secretion into the follicular lumen, Tg is unable to become iodinated and generate T3T4. The biogenesis of Tg requires proper interactions with the protein homeostasis (proteostasis) network to attain its folded state and be secreted.^24^ Extensive co-immunoprecipitation studies have shown that the engagement with proteostasis components differs between misfolding variants and WT, in particular with protein disulfide isomerases (PDIs), lectin chaperones, and HSP70/90 chaperones.^24–28^ Affinity purification mass spectrometry (AP-MS) studies have corroborated the stronger and prolonged interactions of secretion-deficient variants with those proteostasis pathways. AP-MS has also provided a platform to discover previously undetermined interactions.^19,29^

Tg has a large genetic diversity in the human population, with 16 polymorphisms occurring at an allele frequency >1% in Gnomad v4.1.0. More than 160 variants are linked to hypothyroidism, and many more benign variants are reported.^4,30^ This genetic diversity in Tg makes it challenging to determine which sequence variants are causative of hypothyroidism. Previous variants have been primarily discovered through small cohort studies of patients with hypothyroidism.^31–34^ Variants that do not always result in hypothyroidism, such as Q870H, have conflicting pathogenicity assignments.^33,35–37^ Large-scale genetic association studies, on the other hand, allow for more statistical power and a deeper understanding of variable expressivity and disease penetrance for linking Tg variants to thyroid function.^38^ One recently established large-scale biobank is the *All of Us* (AoU) database that collaborates with many institutions within the US, linking genomic data to longitudinal health records, allowing for enhanced genotype to phenotype assessment.^39^

Here, we leveraged the AoU database to link participant medical information on thyroid function to Tg genotype to identify uncharacterized sequence variants that may be causally linked to hypothyroidism. We show that some variants in participants relate to a higher risk of hypothyroidism, while some have a limited effect on thyroid function. Examining variant Tg protein secretion, R152H, R320C, Q870H, and P1494L show a decrease in secretion efficiency in HEK293T and/or Fisher rat thyroid (FRT) cells. Through affinity purification mass spectrometry (AP-MS), we determined how these variants are misprocessed by the proteostasis network. The secretion efficiency for cells that mistrafficked in both HEK293T and FRT cells showed a typical increase in interaction intensity with previously determined proteostasis factors. However, variants A993T and P1494L displayed divergent proteostasis factor engagement, and their occurrence was linked to higher anti-TPO antibody levels in participants, suggesting a divergent hypothyroidism etiology. We estimate the variants characterized here affect ∼150,000 people in the US. In particular, the Q870H variant accounts for ∼100,000 of those individuals. The Q870H variant could be a good target for a therapeutic strategy that may restore Tg secretion to allow the HPT to regulate T3T4 levels instead of LT4 administration.

## Results

### Frequent Tg variants are linked to an increased risk of hypothyroidism

Our first goal was to find less characterized Tg variants that are associated with thyroid dyshormonogenesis. We focused on “common” Tg variants in the general population and looked at specific variants that were previously identified in patients with congenital hypothyroidism.^4^ The allele count, an indicator of variant commonality, was retrieved from the Gnomad database v4.1.0 to prioritize variants (Table S1).^30^ Next, we linked these variants with data from the *All of Us* (AoU) database. AoU collaborates with many institutions in the US with the aim of enrolling individuals from diverse backgrounds to accelerate biomedical research to improve human health. It includes approximately 250,000 participants with genomic data to connect health records to genotypes.^39,40^ In the clinic, TSH levels are used to diagnose hypothyroidism due to the negative feedback in the HPT axis. Therefore, reported TSH levels in AoU should provide a measurement to determine thyroid efficiency associated with each variant.^3,5,9,41^ Importantly, we excluded participants taking LT4 to avoid confounding variables since administration of LT4 reduces TSH levels. We compared the most recent TSH measurement from AoU participants with 10 Tg variants (Fig. 1A). Normal levels of TSH are between 0.5 and 4.5 milli-international units per liter (mlU/L), with lower values associated with hyperthyroidism and elevated values with hypothyroidism.^9^ Some of the variants showed a slight increase in the median TSH levels compared to the variant with the lowest levels of TSH (L1063M), including A993T, P1494L, R320C, Q870H, T1498M, R152H, N2616I, and P1012L (Fig. 1A). However, none of the TSH levels were significantly higher. Meanwhile, D1767del and L1063M had similarly low levels of TSH, indicating these variants likely retain Tg function. A key consideration to these measurements is that all the participants in AoU are heterozygous for the variants indicated, which may dampen a more severe phenotype.

**Figure 1.**
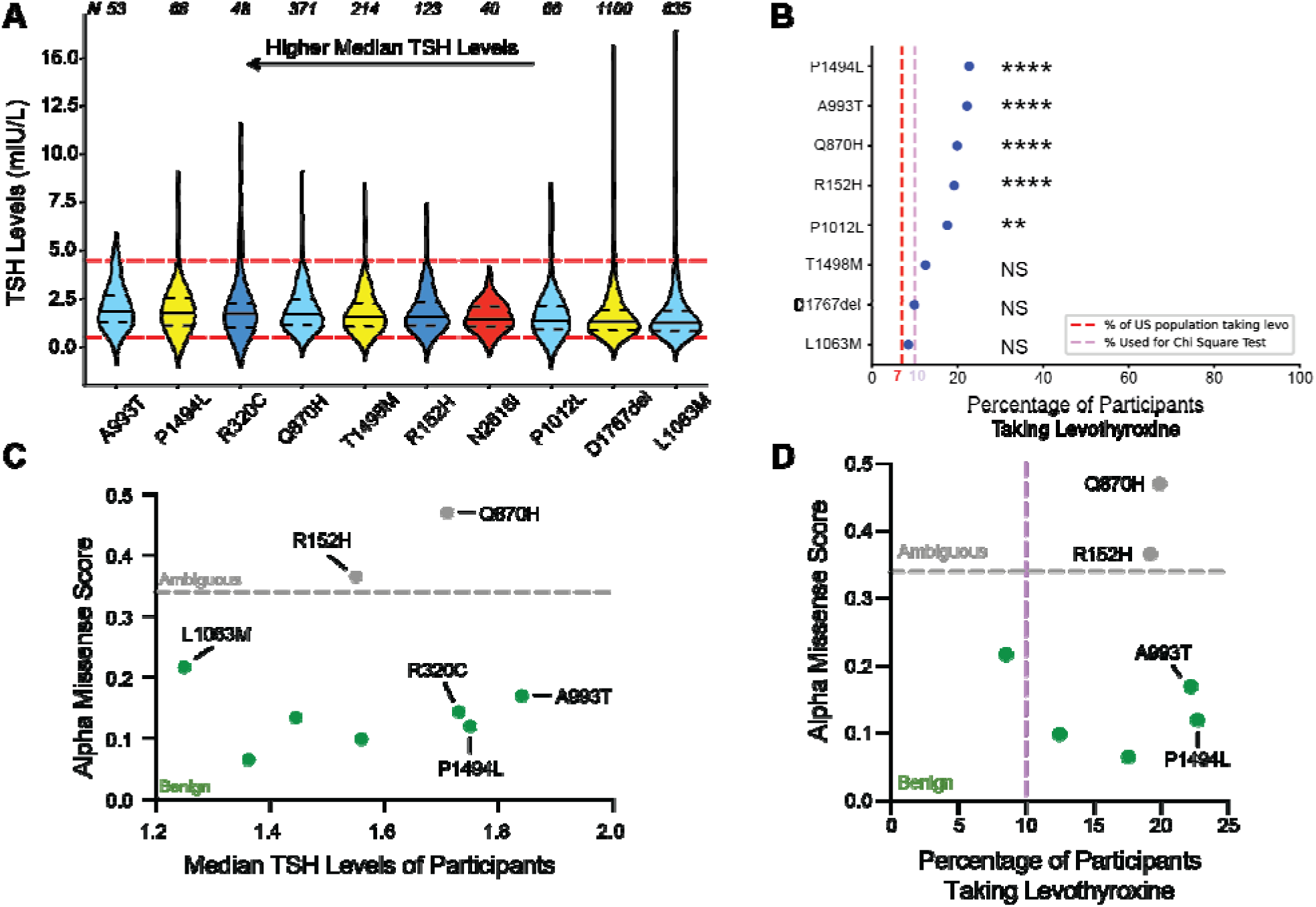
Variants found in AoU participants show a decrease in thyroid function. **A)** Distribution of the most recent measurement of TSH levels for AoU participants who do not take levothyroxine. Variants are sorted from the highest median TSH level to the lowest median TSH level.^9^ Fill color is based on the domain of Tg the variant is present in (see Fig. 2B). Red dashed lines represent the upper and lower limit of normal TSH levels. Lines within each violin represent quartiles. N listed above each violin plot. **B)** Percentage of participants who take levothyroxine. ^2^ was performed with an expectation of 10%, N > 95. **C)** Comparing the Alpha Missense score to the median levels of TSH among participants. **D)** Probability that participants levothyroxine. Colors of variants are based on Alpha Missense predicted pathogenicity. designates benign (<0.33) and ambiguous in grey (≥0.33). ** p <0.005, **** p <0.0001.

Next, we retrieved the percentage of participants taking LT4 for each of the variants as a proxy for hypothyroidism. We found that 5 of the variants – P1494L, A993T, Q870H, R152H, and P1012L – are associated with an increased probability that a participant carrying these variants is currently medicated with LT4 (Fig. 1B). Although approximately 7% of the US population takes LT4, we implemented a conservative 10% cutoff as background for a χ^2^ comparison to reduce false positives and bolster the variants identified as hypothyroidism risk factors.^6^

Finding that some variants showed a significantly higher probability of taking LT4, we sought to estimate: 1) the number of individuals in the US taking LT4 with those variants; and 2) how many of those individuals could benefit from precision medicine. To estimate the number of individuals in the US who are carriers of these variants and currently take LT4, the allele frequency was multiplied by the US population (∼330 million) and then by the disease penetrance (Table 1). Allele frequency can be influenced by biases in database composition. To make a more conservative estimate, the lower reported allele frequency value from either AoU or gnomAD for each variant was used. Disease penetrance was defined as the percentage of participants with the variant who were reported to use levothyroxine (Fig. 1B). To estimate the population that might benefit from a future precision medicine approach, we subtracted the expected background rate of hypothyroidism (10%) to represent cases due to other causes (Table 1). This approach likely underestimates the true target population for two reasons. First, some individuals in the 10% removed could still benefit from a targeted approach. Second, due to a lack of participants who are homozygous for each variant, we cannot estimate thyroid function for homozygous individuals. Regardless, based on these calculations, approximately 150,000 individuals could benefit from targeted interventions, with roughly 100,000 individuals carrying the Q870H variant.

**Table 1.** Estimation of Americans who take LT4 and could benefit from precision medicine. Variants associated with a significant increase in LT4 usage were analyzed to estimate US prevalence and potential precision medicine impact. Allele frequency was multiplied by the US population (330 million) to estimate the number of Americans carrying each variant. This value was then multiplied by disease penetrance to estimate the number currently taking LT4. The “precision medicine targetable” population was calculated as (penetrance – 10%) × carriers, accounting for the background hypothyroidism rate. *Allele frequency is the lower value from AoU or gnomAD.

| Variant | Allele Freq* | Americans with Variant (×10<sup>3</sup>) | % Penetrance | Americans Taking Levothyroxine | Precision Medicine Targetable |
| --- | --- | --- | --- | --- | --- |
| R152H | 0.00066 | 225.42 | 19.2 | 43281 | 20739 |
| Q870H | 0.00297 | 1011.16 | 19.9 | 201221 | 100105 |
| A993T | 0.00032 | 109.82 | 22.2 | 24380 | 13399 |
| P1012L | 0.00071 | 241.06 | 17.6 | 42427 | 18321 |
| P1494L | 0.00056 | 190.40 | 22.7 | 43221 | 24181 |

*Note:*
Allele frequency estimated from lower value between gnomAD and All of Us

We next tested to see if any of these variant pathogenicities can be predicted through computationally predictive methods such as AlphaMissense.^42^ AlphaMissense employs a neural network based on multiple sequence alignments, variant frequency in the human population, and protein structural information to predict pathogenicity of all given missense mutations across the human genome in coding sequences. Q870H and R152H displayed AlphaMissense scores in the ambiguous range (Fig. 1C, D). This result is in line with some participants showing an increased likelihood of taking LT4. Other pathogenic variants, such as A993T, P1012L, and P1494L, are predicted to be benign. Alpha missense scores show no clear correlation between participant metrics of TSH or percentage of participants taking levothyroxine (Fig. 1C, D). Overall, this analysis shows variants found in Tg can have a large disease burden on people but are not predictable as pathogenic by current computational methods. This further highlights the necessity and power of using large-scale biomedical databases to identify genetic susceptibility for hypothyroidism due to Tg variants.

### Structure-mapping finds high-risk Tg variants at residues promoting molecular interactions

Tg is a 330 kDa protein with multiple domains, including Tg repeat, cholinesterase-like (ChEL), arm, and core domains (Fig. 2A).^15^ The ChEL domain acts as a chaperone for the rest of Tg that is otherwise unable to fold properly and get secreted from cells.^43^ Dimerization at the core and the ChEL interface is required for Tg secretion into the follicular lumen and for subsequent T3T4 production.^15,44^ Additionally, there are 60 disulfide bonds present in Tg that help stabilize the structure, and alteration to proper disulfide bond formation can lead to misfolding and diminished secretion.^15^

**Figure 2.**
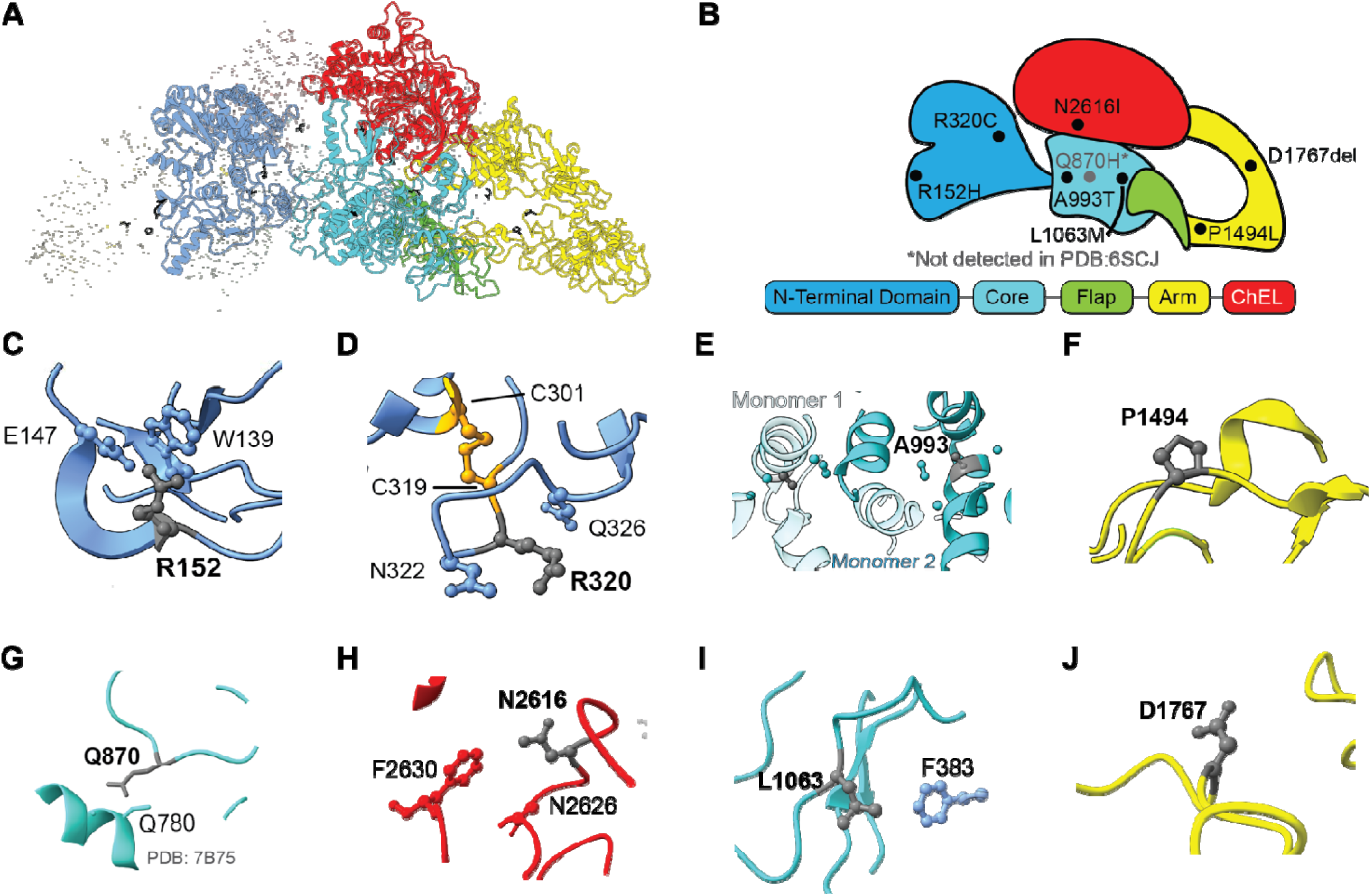
Structural analysis of Tg variants shows stabilizing molecular interactions. **A)** Cryo-EM structure of Tg, where domains are color-coded as shown in the figure legend. Dark blue: N-terminal domain; light blue: core domain; green: flap; yellow: arm; and red: ChEL domain. Variant residues are colored black. PDB: 6SCJ. **B)** Cartoon schematic of thyroglobulin with variant locations represented by black dots. Q870 was not resolved in 6SCJ, but the approximate location is highlighted. **C)** R152 forms a salt bridge with E147 and forms a cation-pi interaction with W139. **D)** R320 can form hydrogen bonds with N322 and Q326, but a disulfide bond between C319 and C301 is one residue away and may be altered with the R320C variant. **E)** A993 is found in a hydrophobic pocket at the dimer interface with the A993 on the other monomer in close proximity. **F)** P1494 does not form interactions with other residues but may add structural rigidity in the loop. **G)** Q870 is not resolved in PDB:6SCJ. Shown is a close-up of PDB:7B75, where it forms a hydrogen bond with Q780. **H)** N2616 can engage in hydrogen bonding with N2616. **I)** L1063 is found in a flexible region and can form van der Waals interactions with F383. **J)** D1767 does not form interactions with nearby residues and is found in a flexible region.

To further understand how the variants could affect Tg structure, we mapped the mutations to previously determined structures of Tg.^15,45^ The variants are located throughout the different domains of Tg (Fig. 2B). R152 forms a salt bridge with the nearby E147 and engages in a cation-π interaction with W139 (Fig 2C). Next, R320C introduces a new cysteine residue, which could lead to incorrect disulfide bond formation. The nearest disulfide bond is just one residue away, occurring between C301 and C319 (Fig. 2D). A993T is found at the dimer interface in a hydrophobic pocket that is separated by an alpha helix relative to the A993 residue in the other monomer (Fig. 2E). P1494L is found within the arm domain and may contribute to entropic stability within the protein structure (Fig. 2F). Interestingly, Q870 is in an unresolved region in PDB:6SCJ, but in a different structure (PDB:7B75), the side chain forms a hydrogen bond with Q780 (Fig. 2G). N2616 engages in hydrogen bonds with N2626 (Fig. 2H). Overall, the variants that appear associated with decreased thyroid efficiency and/or the likelihood that a person is taking LT4 occur in residues with molecular interactions with nearby residues. We next looked at the variants L1063M and D1767del that were associated with the smallest impact on thyroid efficiency (Fig. 1A, B). L1063 is found within a flexible region that forms van der Waals interactions with F383 (Fig. 2I). Meanwhile, D1767 shows no interactions with nearby residues and is found within a flexible region of the protein (Fig. 2J). These two variants are found in more flexible regions, with D1767 showing no apparent molecular interactions with other residues, making these modifications at these two residues less likely to destabilize Tg structure.

### Disease-Associated Tg Variants Show Secretion Defects

Since other known Tg variants show misfolding as a disease mechanism for hypothyroidism, we hypothesized that some of the variants investigated here may exhibit similar misfolding and improper trafficking out of the cell.^18–22^ Therefore, we measured Tg secretion from HEK293T cells transiently transfected with Tg-FLAG constructs. We used C1264R Tg as a control, which is a known disease-causing variant that is only minimally secreted (< 1% of WT levels).^19,29,46,47^ Unlike C1264R, the variants tested were secreted in detectable amounts, but R152H, R320C, and Q870H exhibited a significant reduction in secretion efficiency compared to WT (Fig. 3A, B). Intriguingly, these three variants occurred in people who exhibited higher TSH levels and/or increased risk of taking LT4. A993T and P1494L, which showed the highest probability of LT4 usage, showed a mild decrease in secretion efficiency compared to WT, even though it was not statistically significant.

**Figure 3.**
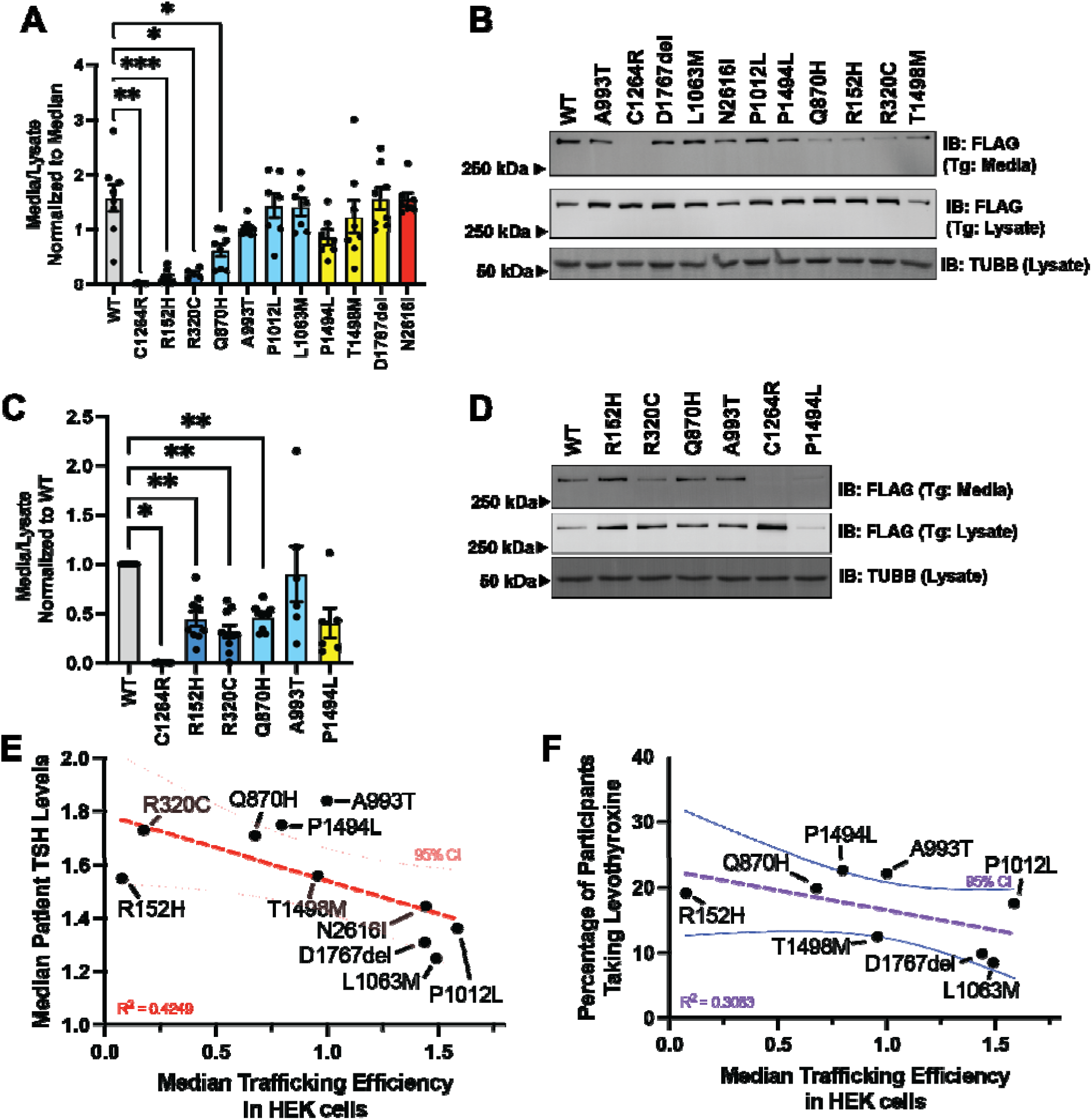
R152H, R320C, and Q870H Tg variants are not fully secreted out of the cell. **A)** Secretion efficiency of Tg in HEK293T cells, measured as the ratio of the Tg detected in the media divided by the Tg detected in the lysate. Data was normalized to the mutant with the median secretion efficiency. One-way ANOVA with Fisher’s LSD correction compared to WT. N ≥ 5 **B)** Representative blot of Tg lysate and media levels in HEK293T cell to assess secretion efficiency shown in **A**. **C)** Secretion efficiency of Tg in FRT cells as measured by Tg levels in the media divided by the Tg levels in cell lysate, normalized to WT. Wilcoxon one-sample T Test, N ≥ 6 **D)** Representative blot of FRT cell secretion shown in **C**. **E)** Comparison of the secretion efficiency of Tg variants in HEK293T to TSH levels in patients. **F)** Comparison of the secretion efficiency to the percentage of participants taking levothyroxine. Bar graph error bars represent the average ± SEM. The dashed line on the graph represents linear regression. * p < 0.05, ** p < 0.01, *** p < 0.001.

Next, we validated the expression and secretion in isogenic Fischer Rat Thyroid (FRT) cells engineered to stably express each Tg construct. We investigated the secretion of three deficient mutants, along with A993T and P1494L, which were associated with a higher percentage of carriers taking LT4 (Fig. 1D). R152H, R320C, and Q870H, which had impaired secretion in HEK293T cells, also showed a reduced secretion efficiency in FRT cells (Fig. 3C, D). The results validate that the secretion deficiency is also present in thyroid-derived cells. Interestingly, P1494L also had decreased secretion efficiency in FRT cells, albeit not statistically significant (p = 0.0625), similar to what was observed in HEK293T cells.

To further assess the relationship between Tg protein secretion and thyroid function, we compared median TSH levels of participants to the median secretion efficiency in HEK cells. These two metrics exhibited a Pearson correlation coefficient (r^2^) of 0.4249, with A993T and P1494L falling outside the 95% confidence interval (Fig. 3E). Comparing secretion efficiency in HEK cells to the percentage of participants taking LT4 resulted in a mild correlation of 0.3083 with A993T, P1494L, and T1498M falling outside the 95% confidence interval (Fig. 3F). Overall, these results are indicative of a general trend that variants with lower secretion efficiency have less functioning thyroids. Nonetheless, A993T and P1494L are outliers in both analyses, suggesting other factors may be important to the etiology of these two variants.

### The Secretion Defect of Disease-associated Tg Variants Can be Attributed to Altered Proteostasis Interactions

Three of the newly characterized variants had secretion deficiencies, thus motivating further investigations into the underlying proteostasis defects to gain insight into how variants are mishandled.^19,29,48,49^ To determine how the different Tg variants interact with the proteostasis network, we employed affinity purification coupled to tandem mass tag-multiplexed mass spectrometry (AP-MS). We used an in situ chemical crosslinker to capture transient interactions between Tg and the proteostasis factors in intact FRT cells that stably express Tg-FLAG constructs. Lysed samples were then co-immunoprecipitated using anti-FLAG beads, followed by elution, trypsin digestion, and tandem mass tag (TMT) labeling, followed by liquid chromatography-tandem mass spectrometry (LC-MS/MS) analysis. We chose to investigate the variants that trafficked poorly in FRT cells (R152H, R320C, and Q870H). In addition, we included variants that were associated with the highest participant percentage taking LT4 (A993T and P1494L). We also used WT and C1264R as a secretion-deficient control for comparisons, as well as FRT cells not expressing Tg (mock) for background filtering (n=8).

The first goal was to determine the enrichment of proteostasis interactors for the six Tg variants. After median normalization of TMT intensity (Fig. S1A, B), we compared the abundance of identified proteins in each affinity purification to the FRT mock condition to determine the co-immunoprecipitated proteins for each Tg variant (Fig. S2A-H, Table S2). We identified 133 total proteins and found moderate overlap (30 interactors) with prior studies on Tg interactomes (Fig. S3).^19,29^ Moderate overlap can be attributed to divergent conditions– one study was performed in HEK293T cells, and the time-resolved FRT cells study incorporated noncanonical amino acid tagging, click chemistry derivatization, and tandem affinity purifications. We generally found variants with better secretion efficiencies had fewer interactors, particularly A993T (Fig. S2B, H). Meanwhile, R152H, R320C, and P1494L had more interactors compared to WT. Q870H showed intermediate interactors, but the most extreme misfolding variant C1264R showed the most interactors (Fig. S2C-H). We then collated the significantly enriched proteins, creating a master list of Tg interactions. These TMT intensities of enriched proteins were normalized to the Tg bait protein to account for slight expression differences, and then log_2_ fold changes (FC) compared to WT were calculated (Table S3). To assess deviations in interaction profiles between the Tg variants, k-means clustering of the log_2_ FC was carried out. Five clusters were determined as optimum based on silhouette and elbow plots. (Fig. S4A, B). This organization revealed clusters of interactors with higher enrichment specificity for secretion-deficient mutants (cluster 1), for WT (cluster 3), for A993T, C1264R, and P1494L (cluster 4), and finally for the secretion-sufficient A993T (cluster 5) (Fig. 4, Fig. S5A-D, Fig. S6). We then mapped protein IDs to distinct proteostasis pathways using the Proteostasis Consortium annotations to understand how the interactions with the proteostasis network change across the variants.^50,51^ Cluster 1 is composed primarily of ER proteostasis factors showing the highest interactions with R152H, R320C, and C1264R (Fig. 4), which exhibited secretion defects in both HEK293T and FRT cells (Fig. 3A-D). This includes interactions with lectin chaperones (Calr and Canx), protein disulfide isomerases (Pdia3/4/6, P4hb, Erp29, and Dnajc10), and other HSP70/90 chaperones (Hspa5 and Hsp90B1). This cluster includes many previously described interactors that are important for folding of Tg, which were shown to have higher engagement with misfolding-prone mutants.^24–28^ Meanwhile, cluster 4 exhibits the highest interactions for A993T, P1494L and C1264R and includes proteins involved in translation (Fig. 4). Cluster 5 shows the strong interactions most exclusively for A993T, which is properly secreted from the cell. The two most represented proteostasis branches in cluster 5 are the ubiquitin proteasome system and ER proteostasis (Fig. 4), which may indicate that A993T is still more prone to misfolding than WT Tg even though this variant is secreted to similar levels.

**Figure 4.**
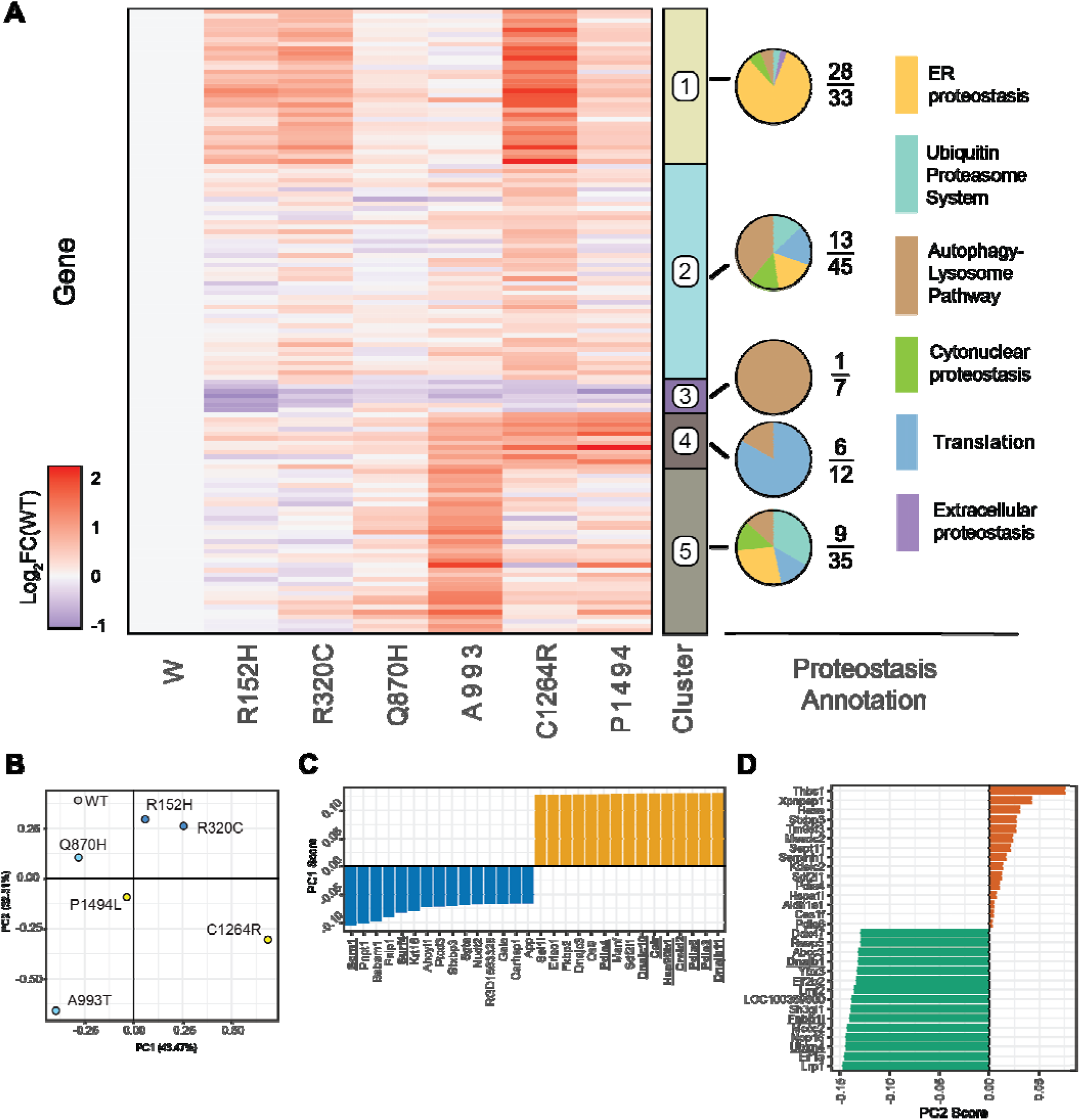
Tg variants show differential interactions with the proteostasis network. **A)** Heatmap of all statistically enriched proteins, showing their relevant enrichment compared to WT (Log_2_FC(WT)). This data was then clustered by k-means to determine unique enrichment profiles between the mutants. Proteins in each cluster were assigned to the Proteostasis Consortium branch classifications.^50,51^ Pie charts (right) represent the percentage of identified proteins and the branch they are associated with. Genes that were not found in the consortium list are not considered in the pie charts. The numerator shows the number of proteins in the given cluster that have annotations and are represented in the pie chart. The denominator represents the total number of proteins in the cluster. **B)** PCA plot based on bait normalized Log_2_FC(WT) using only statistically enriched proteins. **C)** The top 15 positive and negative loading scores for PC1 and **D)** PC2.

Next, we were interested in the global similarity in the interactomes of the Tg variants to determine which variants behave most similarly, and to determine the primary proteostasis factors that drive the similarity. We carried out a principal component analysis (PCA) of interactor log_2_ FC for all the variants (Fig. 4B). This analysis revealed that Q870H shows the most global similarity to WT. The most severe secretion-deficient mutant C1264R is the most distant from WT in both the PC1 and PC2 direction (in opposing quadrants from one another). Lastly, R152H and R320C, which were secretion-deficient in both cell lines, deviate primarily in the positive PC1 direction compared to WT. When assessing the loadings for PC directions, the largest contributing proteins for the positive PC1 include folding chaperones such as Dnajb11, PDIs, Calr and HSP90 (Fig. 4C). Genes associated with more negative loading scores are trafficking factors such as Scrn1 implicated in exocytosis and Surf4 which is responsible for trafficking many proteins out of the ER (Fig 4C).^52–54^ A993T, which had WT-like secretion in both HEK293T and FRT cells, showed minimal movement in the PC1 direction but shifts primarily in the negative PC2 direction compared to WT. High loading scores associated with this PC2 shift include factors associated with the autophagic pathway, Fnbp1l and Dnajb1, and a factor involved in ERAD, Ubxn4. (Fig. 4D). P1494L, which showed mild secretion deficiency in HEK293T cells but was secretion-deficient in FRT cells, also primarily deviated in the negative PC2 direction compared to WT. Overall, the secretion efficiencies in the two different cell lines are correlated with the variant placement along the PC axes of the PCA plot, except for Q870H, which showed a milder defect. The interactome analysis revealed that the Tg secretion defects can be partially attributed to the altered proteostasis engagements.

### A993T and P1494L Variants Show Interactions with Factors that Link to Hashimoto’s Disease

Even though A993T is a variant associated with an increased likelihood of taking LT4, this variant did not exhibit lowered secretion efficiency compared to WT. However, when looking at the interactions with the proteostasis network, A993T showed unique interactions compared to other variants. Fnbp1l is a protein involved in the autophagy pathway, but specifically in xenophagy, which is responsible for detecting foreign pathogens and can lead to peptide display through MHC-II.^55,56^ Even though the protein was only identified in one out of four total MS runs, two independent AP replicates clearly show higher co-immunoprecipitation levels of Fnbp1l with A993T compared to other variants (Fig. 5A). Additional factors in cluster 5 may be involved in downstream MHC display. DNAJB1 is a cofactor with broad client specificity that can target proteins for lysosomal degradation (Fig. S7A). Additionally, Bag6 and Ubxn4, map to the ubiquitin proteosome system, which generates peptides for MHC loading: Bag6 can target proteins for degradation and reduce the MHC levels on the cell surface, which can be affected by insufficient loading of peptides into the MHC binding pocket (Fig. S7B).^57^ Ubxn4 is an adapter protein to recruit valosin-containing protein (VCP) to the ER membrane for targeting proteins for proteosomal degradation(Fig. S7C).^58^

**Figure 5.**
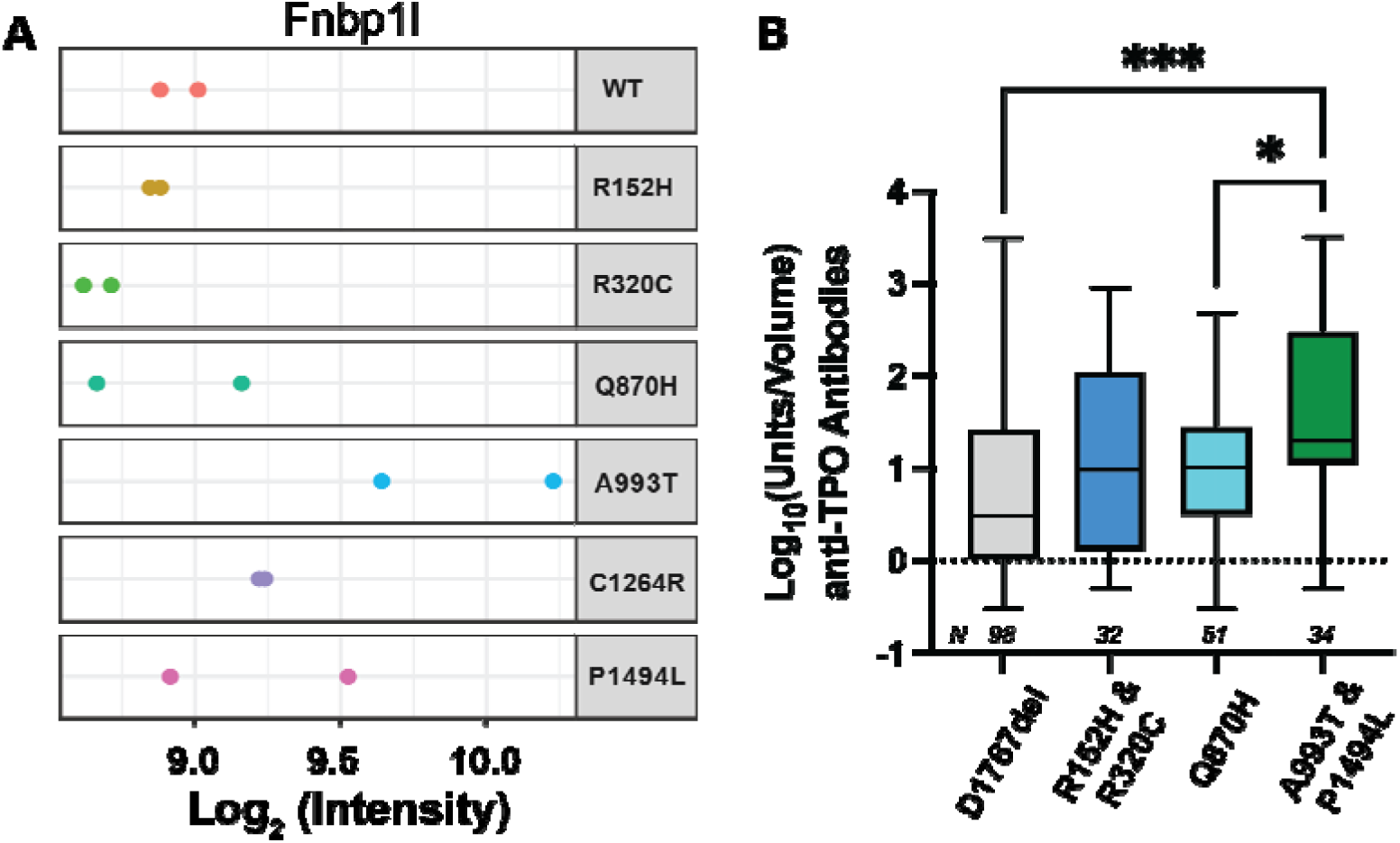
A993T and P1494L Tg variants are associated with factors related to Hashimoto’s disease. **A)** Plot showing the log_2_(intensities) of Fnbp1l detected across different variants. N = 2. **B)** Box plot showing the log_10_(units/volume) of anti-TPO antibodies with mutations bucketed based on quadrants in the PCA plot. One-way ANOVA with Tukey post-hoc. Boxplot lines represent quartiles, and whiskers represent the minimum and maximum. N is listed in the figure. * p < 0.05, *** p < 0.001.

The discovery of autophagy and proteosomal factors as A993T-specific interactors led us to hypothesize that variants shifting negatively in the PC2 direction may lead to an increased risk of developing Hashimoto’s disease. Hashimoto’s is caused by autoantibodies produced against different thyroid components, causing inflammation and decreased thyroid function.^13^ Most cases involve antibodies produced against Tg and TPO. To test whether variants A993T and P1494L might be linked to Hashimoto’s, we returned to the AoU database to determine whether participants carrying the variants have elevated anti-Tg and anti-TPO antibodies. ^59–61^ These antibody screens are less commonly administered than TSH measurements. Because of the infrequent testing and reporting in AoU, we were unable to compare anti-Tg antibodies among the variants. In contrast, anti-TPO antibody tests are more commonly administered and reported in AoU. Nonetheless, we had to reduce granularity by combining two variants found in each quadrant of the PCA plot to reach sufficient population sizes (n ≥ 20) to test for changes (Fig. 5B). Additionally, we included D1767del as a negative control because we found limited effect on thyroid function and Tg secretion efficiency in cell models (Fig 1A, B, Fig 2A-D). Anti-TPO antibody levels exhibited a significant increase for the combined carriers of A993T and P1494L, suggesting variants that shift negatively in the PC2 direction could be at risk of developing Hashimoto’s disease (Fig. 5B). Intriguingly, this is the only group of variants that showed significantly elevated TPO antibody levels. (Fig. 5B). These results suggest that the interactome remodeling of specific Tg variants, along with data on sustained secretion efficiency, can provide an indication of which variants may have increased susceptibility to Hashimoto’s disease.

## Discussion

We show here the utility of the large biomedical cohort research hub *All of Us* (AoU) to identify prevalent variants of Tg and assess how they contribute to hypothyroidism. AoU allows for investigation of variants with conflicting interpretations, such as the Q870H variant, which was found in patients with hyperthyroidism, hypothyroidism, and non-endemic goiter.^33,35–37^ This also allows for detecting differences in susceptibility to disease, as we found variants showed varying degrees of thyroid function and LT4 usage. The variants highlighted in this study have been detected in patients with hypothyroidism, but there is limited molecular characterization. Hence, using AoU could help prioritize variants in the future.^4^

To investigate the molecular consequence of the mutations in Tg, we first examined secretion efficiency, a characteristic found to decrease in many disease-causing variants. This metric correlated with most mutations, consistent with previous findings suggesting that Tg misfolding represents a common pathogenic mechanism underlying variant-associated hypothyroidism risk. Notably, A993T exhibited secretion efficiency similar to WT despite causing decreased thyroid function. This prompted further investigations into proteostasis network interactions that might delineate the protein quality control deficiencies leading to reduced secretion or aberrant protein processing. All disease-causing mutations exhibited divergent interactomes from WT. Detailed analysis of interaction patterns revealed R152H and R320C engaged with previously established ER proteostasis factors, including lectin chaperones, PDIs, and heat shock proteins, consistent with a protein folding defect as the cause for the section deficiency. In contrast, interactions with the proteostasis network diverged further for the A993T and P1494L variants. In particular, A993T clustered separately from the other variants and exhibited unique interactors linked to protein degradation and antigen presentation, suggesting a divergent etiology.

Cellular protein degradation through proteosomal and lysosomal pathways generates peptides for loading onto MHC molecules for cell surface presentation and immune recognition. Our interacome analysis revealed that several proteins involved in these degradation processes, Fnbpl1, Dnajb1, Bag6, and Ubxn4, showed specific interactions with the A993T variant.^55–58^ This altered interaction profile suggests that A993T could undergo aberrant processing, potentially generating novel autoantigens to elicit Hashimoto’s disease. The AoU database enabled further investigation of this potential disease association, again highlighting the utility and the versatility of large-scale biomedical datasets. Using anti-TPO antibodies as a diagnostic proxy for Hashimoto’s disease susceptibility, we found that changes in the specific proteostasis network interactions correlated with antibody levels. Previous reports linked other Tg variants (R2510Q, S734A, and M1027V) to an increased risk for developing Hashimoto’s disease, making these variants promising candidates for future interactome studies.^62,63^ Additionally, endoplasmic reticulum stress has been postulated as an initiating factor for Hashimoto’s pathogenesis, highlighting a further connection to proteostasis disruptions. While Hashimoto’s disease is primarily characterized as an immune system disorder, our data here suggest that specific Tg variants may contribute genetically by altering protein processing pathways that increase susceptibility to antigen presentation and subsequent autoimmune responses.^13,64^

Several limitations should be considered when interpreting our findings. First, we considered participant data in AoU for a given Tg variant regardless of zygosity, without accounting for the other variants that may influence phenotype. No participants were analyzed as homozygous for the variants of interest. Second, while we examined several Tg-related metrics that correlate with thyroid function, thyroid dysfunction is a complex condition likely resulting from multiple contributing factors. Hence, other aspects of thyroid biology may play important roles in the disease etiology, even when these variants are present. Third, our cell-based models are an imprecise representation of the complex thyroid microenvironment. Future studies would benefit from animal models to validate the functional consequences of the identified variants associated with decreased thyroid function. Finally, although we discovered a possible link between A993T and P1494L variants to Hashimoto’s disease, the underlying pathogenic mechanisms remain unclear. Mechanistic studies examining variant-specific peptide presentation, larger cohorts enriched for these variants, and targeted studies could provide greater resolution to determine the relative disease risk conferred by each of these variants to develop Hashimoto’s disease.

Overall, the biobank-driven approach for identifying functionally relevant genetic variants of interest represents a scalable framework applicable to other disease-relevant proteins and biological pathways important for human health. Large-scale biomedical data repositories like AoU can be used to delineate diseases with heterogeneous etiologies, thereby identifying precision medicine approaches that can optimize patient-specific treatments. Our findings illustrate this etiological diversity: misfolding-prone Tg variants may require different therapeutic strategies than variants that predispose to autoimmune thyroiditis. Among the variants characterized here, Q870H emerges as the most promising candidate to target. This misfolding variant affects a substantial population in the United States, while retaining partial trafficking capacity and demonstrating limited disruptions in the proteostasis network interaction. These characteristics suggest that Q870H-associated Tg misfolding may be amenable to correction through pharmacological chaperones – small molecules that bind to and stabilize misfolded proteins.^65^ This therapeutic approach has shown success in other protein misfolding disorders, particularly cystic fibrosis.^65^ Successful pharmacological chaperone rescue of misfolded Tg could restore normal HPT function in Q870H carriers, potentially improving quality of life and providing an alternative therapeutic option to patients who experience persistent symptoms despite standard LT4 replacement therapy.

## Materials and Methods

### All of Us Cohort

This study used data from the *All of Us* Research Program’s Controlled Tier Data set v7, available to authorized users on the Researcher Workbench. Participants were selected for Tg SNPs that resulted in the variants R152H, R320C, Q870H, A993T, P1012L, L1063M, P1494L, T1498M, D1767del, and N2616I. For analysis of TSH levels, participants were only considered if they did not take levothyroxine. To use only one data point per participant, we used the most recent TSH measurement. For anti-TPO measurements, all participants were used, and all measurements of participants were used.

### Plasmid Production

The Tg-FLAG in pcDNA5/FRT used in this study was previously made.^19^ To generate the mutant constructs, site-directed mutagenesis was performed using a Q5 polymerase (New England BioLabs). All nucleotide sequences used for site-directed mutagenesis can be found in Table S4.

### Cell Culture/Engineering

HEK293T cells were cultured in Dulbecco’s modification of Eagle’s medium (DMEM, Corning) supplemented with 10% fetal bovine serum (FBS, Gibco), 1% L-glutamine (200 mM, Gibco), and 1% penicillin/streptomycin (10,000 U; 10,000□μg/ml, Gibco).

FRT cells were cultured in Ham’s F12, Coon’s Modification (F12) media (Gibco, cat. No. 11765054) supplemented with 5% fetal bovine serum (FBS), and 1% penicillin (10,000 U)/streptomycin (10,000 μg/mL). To generate FRT flp-Tg-FT cells, 5□×□10^5^ cells cultured for 1 day were cotransfected with 2.25□μg of flp recombinase pOG44 plasmid and 0.25□μg of Tg-HT pcDNA5 plasmid using Lipofectamine 3000. Cells were then cultured in media containing Hygromycin B (100□μg/mL) for two weeks to select site-specific recombinants.

### Secretion of Tg-FT from HEK293T Cells

Cells were transiently transfected 24 hours after seeding 2 × 10^5^ cells/mL into six well dishes with Tg expression plasmids of respective variants using the calcium phosphate method.^66^ About 18 hours after transfection, new media was placed onto the cells. Cell culture media was collected at the time of harvest to measure secreted Tg. Cells were then washed with cold phosphate-buffered saline (PBS). Lysates were prepared by lysing in Radioimmunoprecipitation assay (RIPA) buffer (50 mM Tris pH 7.5, 150 mM NaCl, 0.1% SDS, 1% Triton X-100, 0.5% deoxycholate) with protease inhibitor cocktail (Roche, 4693159001). Lysis was performed on plate at 4 °C with gentle rocking for 30 minutes. Lysate was collected and cellular debris was cleared by centrifuging at 14,000 x rpm for 10 min at 4 °C. Protein concentration was normalized to the lowest concentration using a Bicinchoninic Acid (BCA) assay (Thermo Scientific, 23225).

### Secretion of Tg-FT from FRT Cells

Engineered FRT cells stably expressing Tg-FLAG constructs were seeded so that cells were >80% confluent ∼48 hours later when harvesting. Media was collected at the time of harvest. Cells were then washed with cold PBS. Lysis with RIPA and protease inhibitor cocktail occurred on plate at 4 °C with gentle rocking for 30 minutes. Lysate was collected, and cellular debris was then cleared by centrifuging at 14,000 x rpm for 10 min at 4 °C. Protein concentration was normalized to the lowest concentration using a BCA assay (Thermo Scientific, 23225).

### Immunoblotting and SDS-PAGE

Normalized lysates were denatured with 1× Laemmli buffer + 100□ mM dithiothreitol (DTT) and heated at 95□°C for 5□min before being separated by 8% Tris-glycine SDS-PAGE. Samples were transferred onto polyvinylidene difluoride (PVDF) membranes (Millipore, IPFL00010) for immunoblotting and blocked using 5% nonfat dry milk dissolved in tris-buffered saline with 0.1% Tween-20 (Fisher, BP337-100) (TBS-T). Primary antibodies were incubated either at room temperature for 2□hr or overnight (∼16 hr) at 4□°C. Membranes were then washed three times with TBS-T for 5 minutes with gentle rocking and incubated at room temperature for 40 minutes in 5% nonfat dry milk dissolved in TBS-T with secondary antibody. Membranes were washed three times with TBS-T and then imaged using a ChemiDoc MP Imaging System (Bio-Rad). Primary antibodies were acquired from commercial sources and used at the indicated dilutions in immunoblotting buffer (5% bovine serum albumin (BSA) in Tris-buffered saline pH 7.5, 0.1% Tween-20, and 0.1% sodium azide): thyroglobulin (1:1000; Proteintech, 21714-1-AP), M2 FLAG (1:1000; Sigma-Aldrich, F1804). Tubulin-Rhodamine primary antibody (BioRad, 12004165) was used at 1:20000 dilution in 5% milk in TBS-T. Secondary antibodies were obtained from commercial sources and used at the indicated dilutions in 5% milk in TBS-T: Goat anti-mouse Star-bright700 (1:10,000; BioRad 12004158), Goat anti-rabbit IRDye800 (1:10,000; LiCor 926-32211).

### Co-immunoprecipitation

Engineered FRT cells were seeded in 10 cm dishes so that on the day of harvest, ∼72 hours after seeding, the cell plates were >80% confluent. Cells were washed 2 times with 5 mL of PBS, followed by adding 5 mL of PBS + 500 μM DSP (Thermo Scientific, 22585) and incubated at 37 °C for 10 min. An additional 5 mL of 200 mM Tris [pH 7.5] was then added to the cells and incubated at 37 °C for 5 min. Cells were then washed with cold PBS, and 500 μL of RIPA and protease inhibitor was added to the plates. Plates were incubated at 4 °C for 30 minutes with gentle rocking. Cell plates were then scraped, and debris was cleared by centrifuging at 14,000 x rpm for 10 min at 4 °C. Protein concentration was normalized to 600 μg of protein in 600 μL using a BCA assay. Normalized lysates were then mixed end over end for 1 hour at 4 °C with 50 μL of Sepharose 4B bead slurry (MilliPore, 4B200) prewashed with PBS. Samples were then centrifuged at 4 °C for 5 min at 300 x g, and the supernatant was collected and transferred to a new tube with 20 μL of pre-equilibrated Pierce anti-FLAG magnetic beads (Thermo Scientific, A36797) and allowed to mix end over end for ∼18 hours. The samples were then washed 4 times with 500 μL of RIPA, discarding the supernatant each time. The protein was then eluted by adding 100 μL of elution buffer (4% SDS, 100 mM Tris [pH 6.8]) and heating at 95 °C for 5 min.

### Mass Spectrometry Sample Preparation

Co-Immunoprecipitation elution was prepared for MS using a single pot, solid phase-enhanced sample preparation (SP3) beads (Cytiva, 65152105050250 and 45152105050250) sample processing method previously described that utilizes a Biomek i5.^67^. Briefly, proteins were first reduced with 5 mM DTT at 60 °C/700 rpm for 30 min, then alkylated with 20 mM IAA for 30 min at 21 °C in the dark. Finally, the reaction was quenched with a second addition of 5 mM DTT for 15 min at 21 °C. This protein was then digested on 8 μL of total SP3 beads (1:1 hydrophobic to hydrophilic) in 50 μL of 50 mM HEPES [pH 8] with 0.5 μg of Trypsin/Lys-C (Promega, V5073) for 10 hours at 37 °C while shaking at 700 rpm. The sample was kept at 4 °C until further use. The supernatant was transferred to a new tube, and 10 μL of water was added. Next, 40 μL of TMTpro reagent resuspended in acetonitrile (Thermo, A44520) was added to the sample and allowed to incubate for 1 hour. After this, 4 μL of freshly made 10% ammonium bicarbonate was added and incubated at room temperature for 1 hour to quench excess TMT reagent. Samples were then pooled and acidified to a pH of less than 2 with formic acid. Sample volume was then reduced to ∼800 μL and then brought back up to ∼1600 μL with buffer A (94.9% H2O, 5% acetonitrile, 0.1% formic acid).

### Liquid Chromatography-Tandem Mass Spectrometry

LC-MS/MS data-dependent analysis was performed using an Exploris480 mass spectrometer (Thermo Scientific) equipped with a Dionex Ultimate 3000 RSLCnano system (Thermo Scientific). MudPIT columns were made as previously described with 600 μL of labeled peptides loaded onto the trap column using a high-pressure chamber. ^68^ To elute peptides from the first C18 phase to the SCX resin of the MudPIT column, 10 μL of Buffer A was injected and the following 90 minute gradient was utilized: 4% B (99.9% acetonitrile, 0.1% formic acid v/v) (5 min hold) ramped to a mobile phase concentration of 40% B over 90 minutes, ramped to 80% B over 5 mins, held at 80% B for 5 mins, then returned to 4% B for 5 mins and then held at 4% B for the remainder of the analysis at a constant flow rate 500 nl/min. Then to fractionate the sample, 10 µL sequential injections of 10, 30, 60, and 100% buffer C (500 mM ammonium acetate in buffer A) in Buffer A, with a final fraction injection of 90% buffer C and 10% buffer B were utilized with a corresponding 130 min gradient at a flow rate of 500 nl/min (4% B for 10 mins then ramped to 40% B over 90 mins, increased to 80% B over 5 mins then held at 80% B for 5 mins and returned to 4% in 5 mins, and held at 4% for the remainder of the analysis). A 3 sec duty cycle was utilized, consisting of a full scan (400-1600 m/z, 120,000 resolution) and subsequent MS/MS spectra collected in TopSpeed acquisition mode. For MS1 scans, the maximum injection time was set to 50 msec with a normalized AGC target of 100%. Ions were selected for MS/MS fragmentation based on the following criteria: MS1 intensity above 1e4, charge state between 2-6, and monoisotopic peak determination set to peptide. Additionally, a dynamic exclusion time of 45 secs was utilized (determined from peptide elution profiles) with a mass tolerance of +/-10 ppm to maximize peptide identifications. MS/MS spectra were collected with a normalized HCD collision energy of 36, 0.4 m/z isolation window, auto-selected for maximum injection time mode, a normalized AGC target of 100%, at a MS2 Orbitrap resolution of 45,000 with a defined first mass of 110 m/z to ensure measurement of TMTPro reporter ions.

### Mass Spectrometry Data Analysis

Proteome Discoverer 2.4 (Thermo Fisher Scientific) was utilized to obtain peptide identifications and TMT-based protein quantification. MS/MS spectra were searched using SEQUEST-HT against a Uniprot SwissProt rat proteome with human Tg manually added (downloaded Aug 27^th^ 2020), contaminant FASTA50 (containing 379 entries), and a decoy database of reversed peptide sequences. Peptide precursor ion mass tolerance was set to 20 ppm, and fragment ion mass tolerance was fixed at 0.02 Da. Only peptides with a minimum length of six amino acids were considered, and trypsin digestion was assumed with a maximum of two missed cleavages allowed. Dynamic modifications included oxidation of methionine (+15.995 Da), protein N-terminal methionine loss (-131.040 Da), protein N-terminal acetylation (+42.011 Da), and a combination of N-terminal methionine loss and acetylation (-89.030 Da). Static modifications were applied for cysteine carbamidomethylation (+57.021 Da) and TMTpro labeling on the N-terminus or lysines (+304.207 Da). To control false discoveries, the peptide-level false discovery rate (FDR) was set to 1% using the Percolator algorithm. Quantification of TMT reporter ions was performed by including only those with an average signal-to-noise ratio greater than 10:1 and a co-isolation percentage of less than 25%. TMT reporter ion intensities were summed for peptides assigned to the same protein, including razor peptides, which are shared by multiple proteins but assigned to the one providing the most confident identification. Protein identifications were filtered at a 1% FDR threshold, and protein grouping was conducted according to the parsimony principle, which minimizes the number of protein groups assigned to each peptide. Analysis was subsequently carried out using custom R code available on GitHub (https://github.com/Plate-Lab/). Briefly, median normalization was performed across TMT channels for statistical analysis and determination of purified proteins. Downstream, intensities were bait normalized to Tg within each variant.

### Statistical Analysis

For determining the increased likelihood for participants with variants to take LT4, a χ^2^ test was used (Fig. 1). While 7% of the US population takes levothyroxine, we used 10% as the cutoff to reduce false positives and bolster results. Only variants in which at least 20 participants were found to either take levothyroxine or not take levothyroxine for each condition were considered. For anti-TPO antibody levels, a single outlier was removed using a Grubb’s test with a q value of <1.0. For western blot secretion data in both HEK293T and FRT cells, outliers were removed using a Grubbs test with a q value <1.0. This resulted in one outlier being removed for both P1012L and L1063M in HEK293T cell data and one replicate for R152H in the FRT cell data. HEK293T cell secretion of variant significance was determined through a one-way ANOVA with Fisher’s LSD correction comparing all variants to the secretion of WT (Fig. 2). Due to variability in WT secretion for HEK293T cell data, we normalized to the median of all the mutations. Significance of FRT secretion was determined from a Wilcoxon one-sample T test that compared WT (normalized value of 1) to the variants (Fig. 2). When determining significantly enriched proteins in co-immunoprecipitation experiments, we used a T test (Fig. S2). An outlier from anti-TPO measurements was removed, followed by a one-way ANOVA with Dunnett post-hoc comparing all variant groups to L1063M and D1767del (Fig. 5).

## Data, Materials, and Software Availability

Mass spectrometry data have been deposited to the ProteomeXchange Consortium via the PRIDE partner repository with the project accession (PXD069688). The reviewers can access the data using the account information below:

Username

Password: vcX8xLsdTsqD

## Author Contributions

Designed research: J.N.H, L.P.; Performed research: J.N.H, A.D.H. ; Contributed new reagents/analytic tools: J.N.H.; Analyzed data: J.N.H, A.D.H., L.P ; Wrote the paper: J.N.H, L.P.

## Acknowledgements

We gratefully acknowledge *All of Us* participants for their contributions, without whom this research would not have been possible. We also thank the National Institutes of Health’s *All of Us* Research Program for making available the participant data, samples, and cohort examined in this study. We acknowledge funding from the National Institutes of Health: grant R35GM133552 (NIGMS) and grant R01NS095989 (NINDS).

