## Supporting Information for "Public Cohort Analysis Identifies Thyroglobulin Variants as Hypothyroidism Risk Factors"

Jake N Hermanson[1], Andrew D Hudson[2], Lars Plate*[1,2,3]
[1] Department of Biological Sciences, Vanderbilt University, Nashville, Tennessee
[2] Department of Chemistry, Vanderbilt University Nashville, Tennessee
[3] Department of Pathology, Microbiology and Immunology, Vanderbilt University Medical Center, Nashville, Tennessee

### Supplemental Figures.

1. Median normalization of TMT intensities
2. Determining statistically enriched interactors of Tg
3. Venn diagram comparing Tg interactors to previous studies
4. Determining number of clusters to use in k-means algorithm
5. Individual cluster heat maps for clusters 1 to 4
6. Individual cluster heat map for cluster 5
7. Highlights of genes that may result in increased probability to Hashimoto's

**A)**

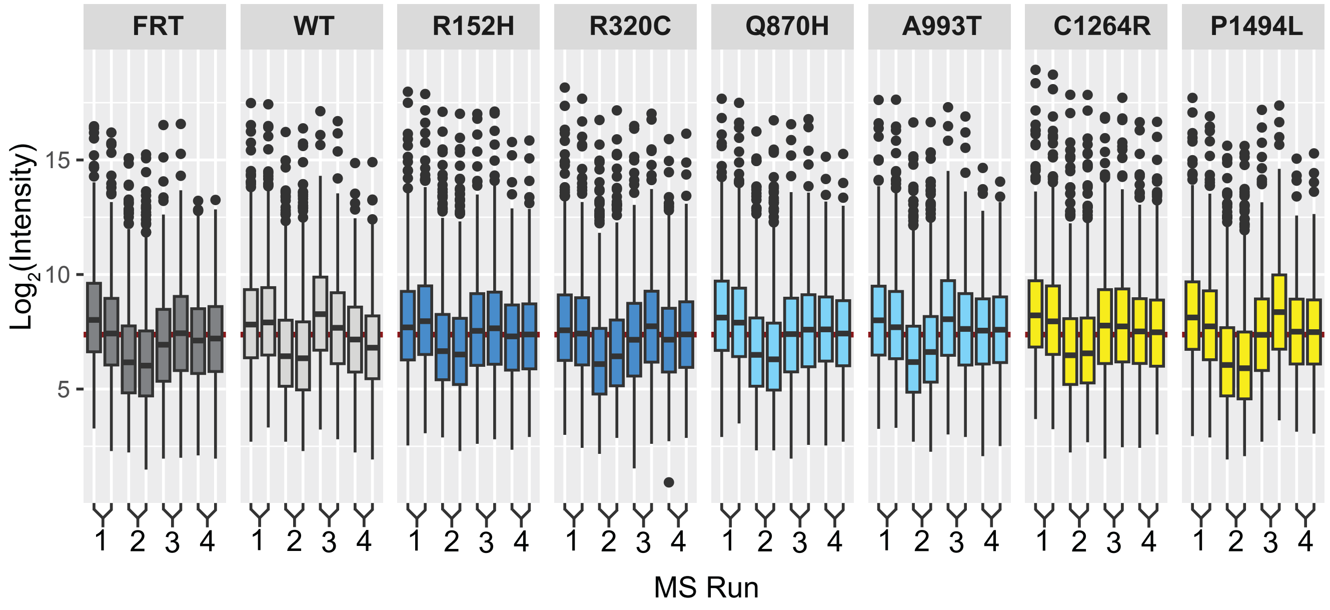


**B)**

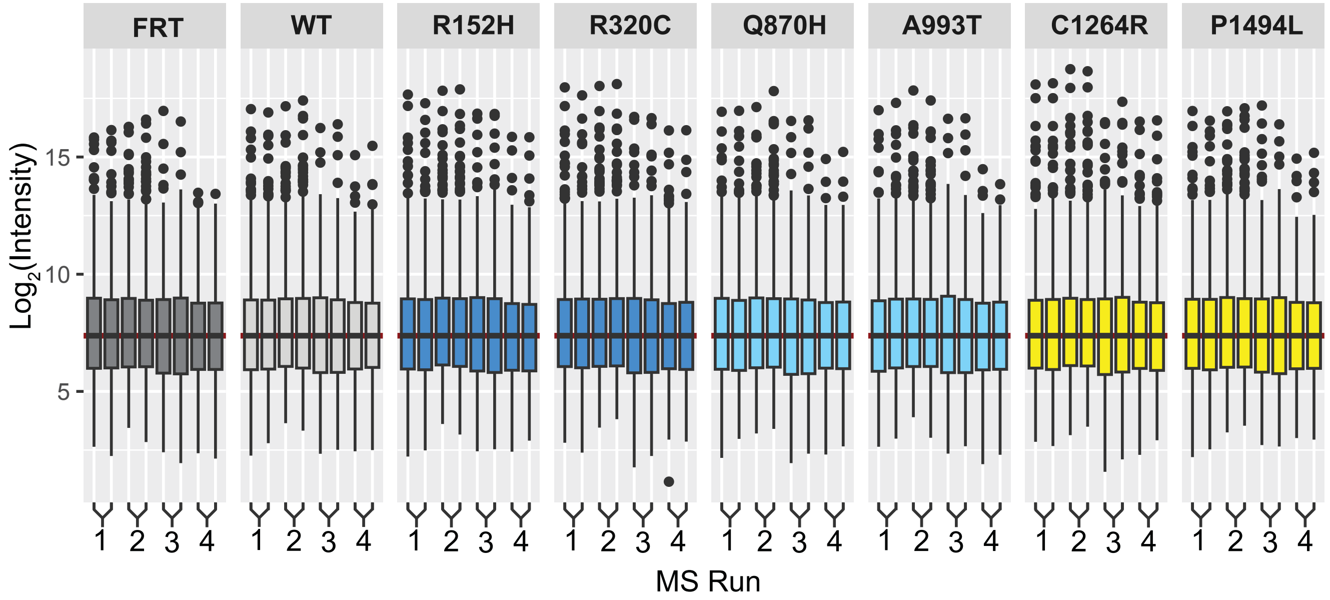


**Figure S1. Median normalization of TMT intensities.** TMT intensities for all protein intensities across MS runs a both **A)** pre and **B)** post global median normalization.

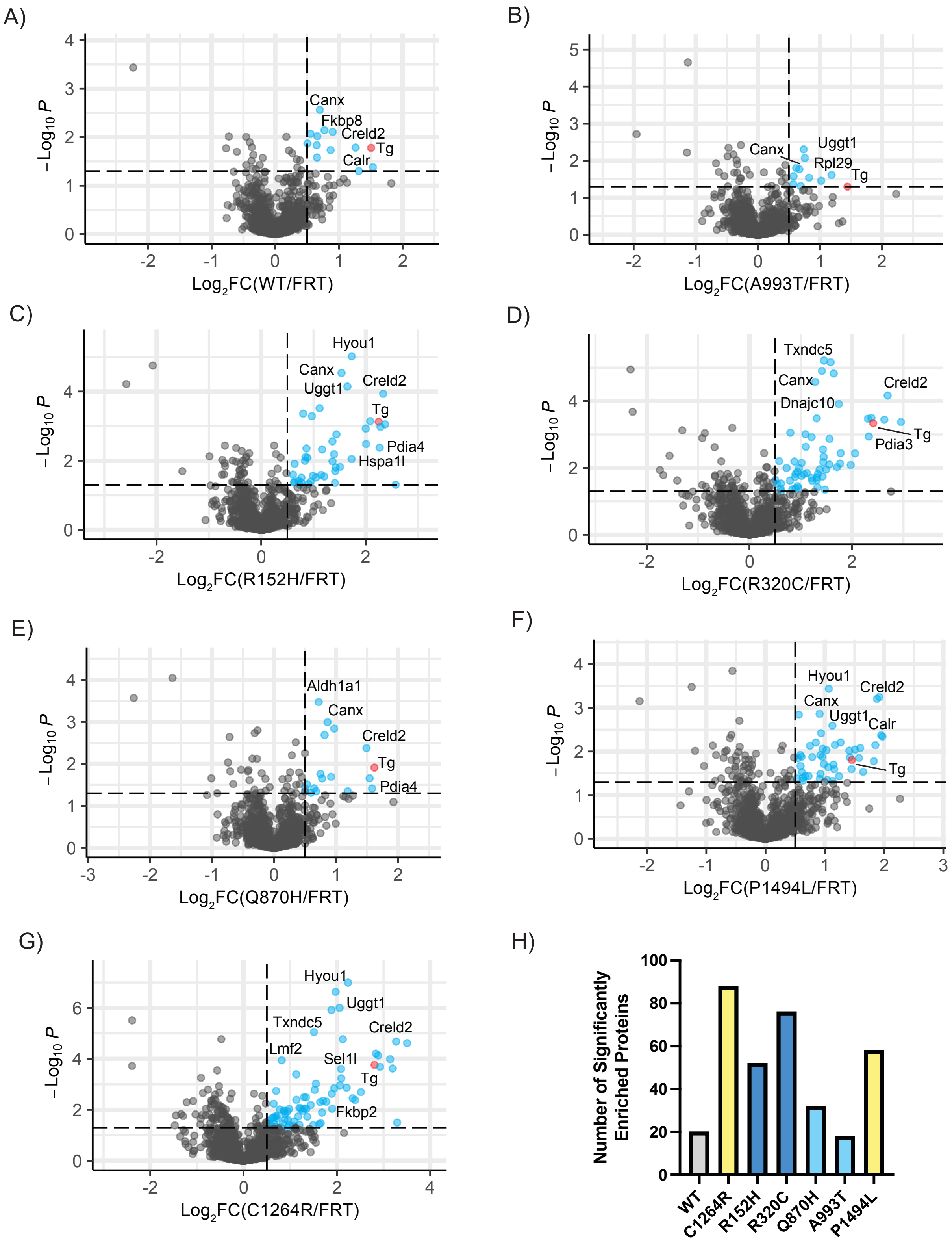


**Figure S2. Determining statistically enriched interactors of Tg.** Volcano plots where the vertical dashed line is the fold change cut off equal to 0.5 and the horizontal dashed line is equal to a -Log$_{10}$P of ~1.3 (p value of 0.05). Cyan dots represent proteins that meet both criteria and the bait protein, Tg, is highlighted red. This was done compared to FRT cell lines not expressing Tg and compared to **A)** WT, **B)** A993T, **C)** R152H, **D)** R320C, **E)** Q870H, **F)** P1494L and **G)** C1264R. T-test, N = 8. **H)** Number of statistically significant interactors determined by MS for each variant.

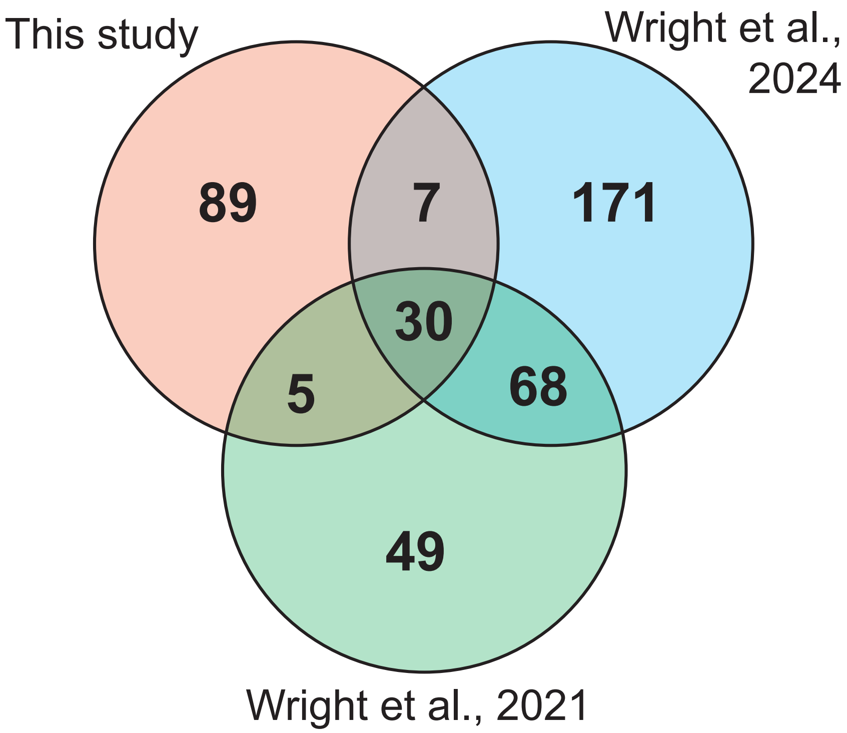


**Figure S3 Comparing Tg interactors to previous studies.** Overlap of statistically significant interactors visualized with a Venn diagram. Red represents interactors found in this study, blue is Wright et al., 2024 which was performed using noncanonical amino acid incorporation in FRT cells.[1] Green represents interactors found in HEK293T cells in Wright et al., 2021.[2]

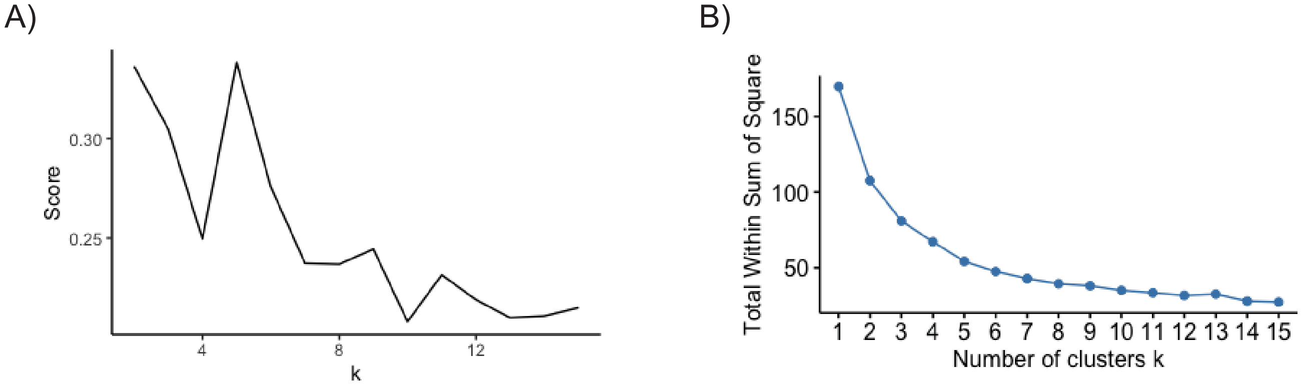


**Figure S4. Determining optimal number of clusters for k-means clustering. A)** Silhouette plot and **B)** Elbow plot.

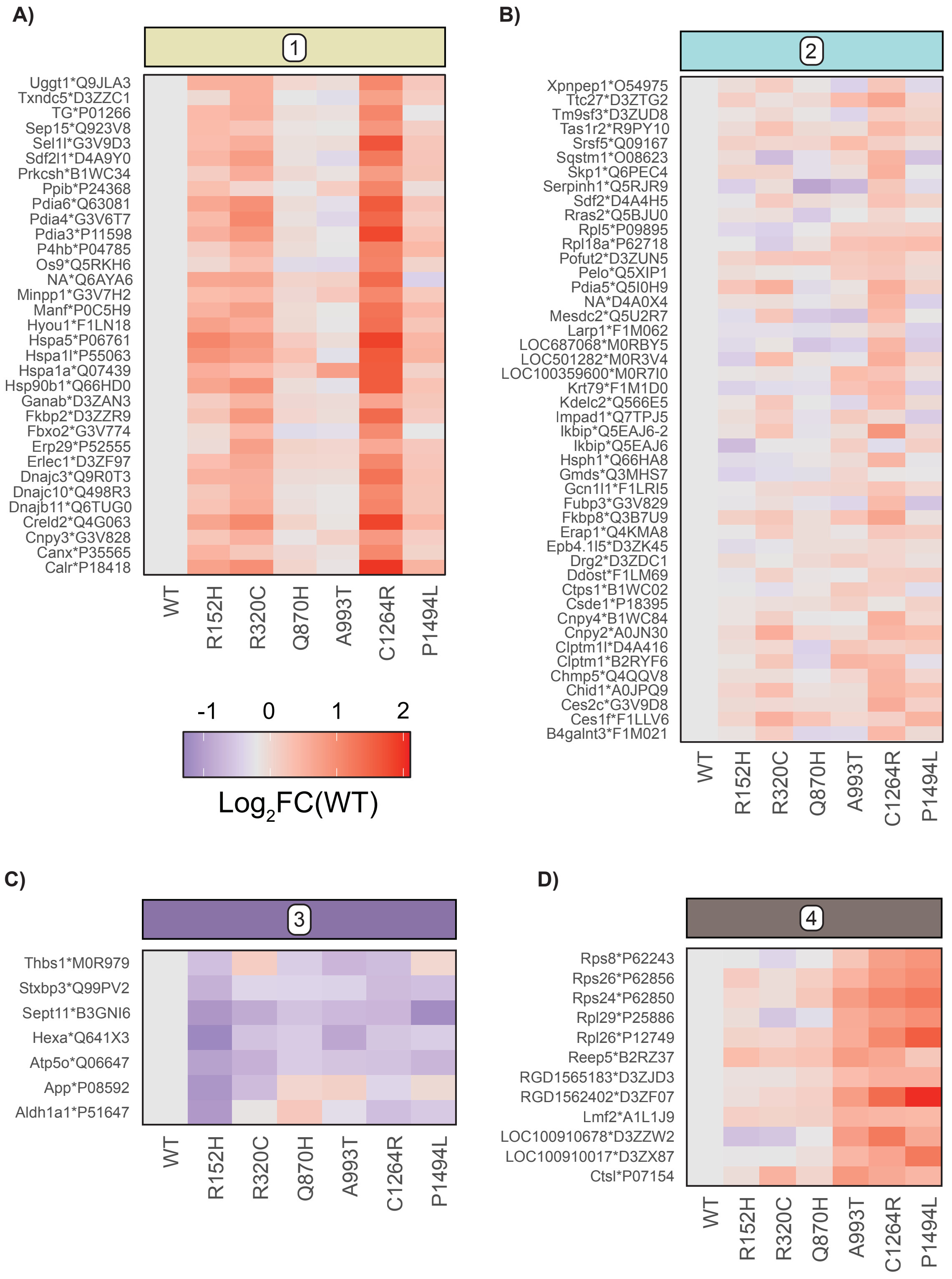


**Figure S5. Individual heat maps of clusters 1 to 4.** Individual genes shown are labelled in ascending order of gene symbol. The annotation is gene*accession where gene is the gene symbol and accession corresponds to the Uniprot accession numbers. This is done for clusters **A)** 1, **B)** 2, **C)** 3 and **D)** 4.

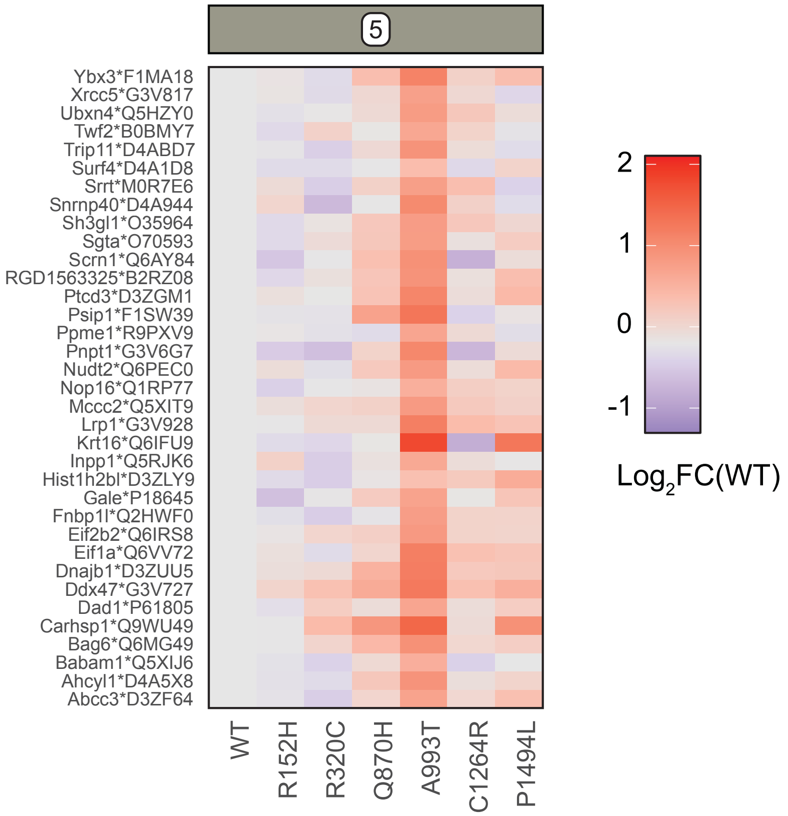


**Figure S6. Individual cluster heatmap of cluster 5.** Individual genes shown are labelled in ascending order of gene symbol. The annotation is gene*accession where gene is the gene symbol and accession are the uniprot accession number.

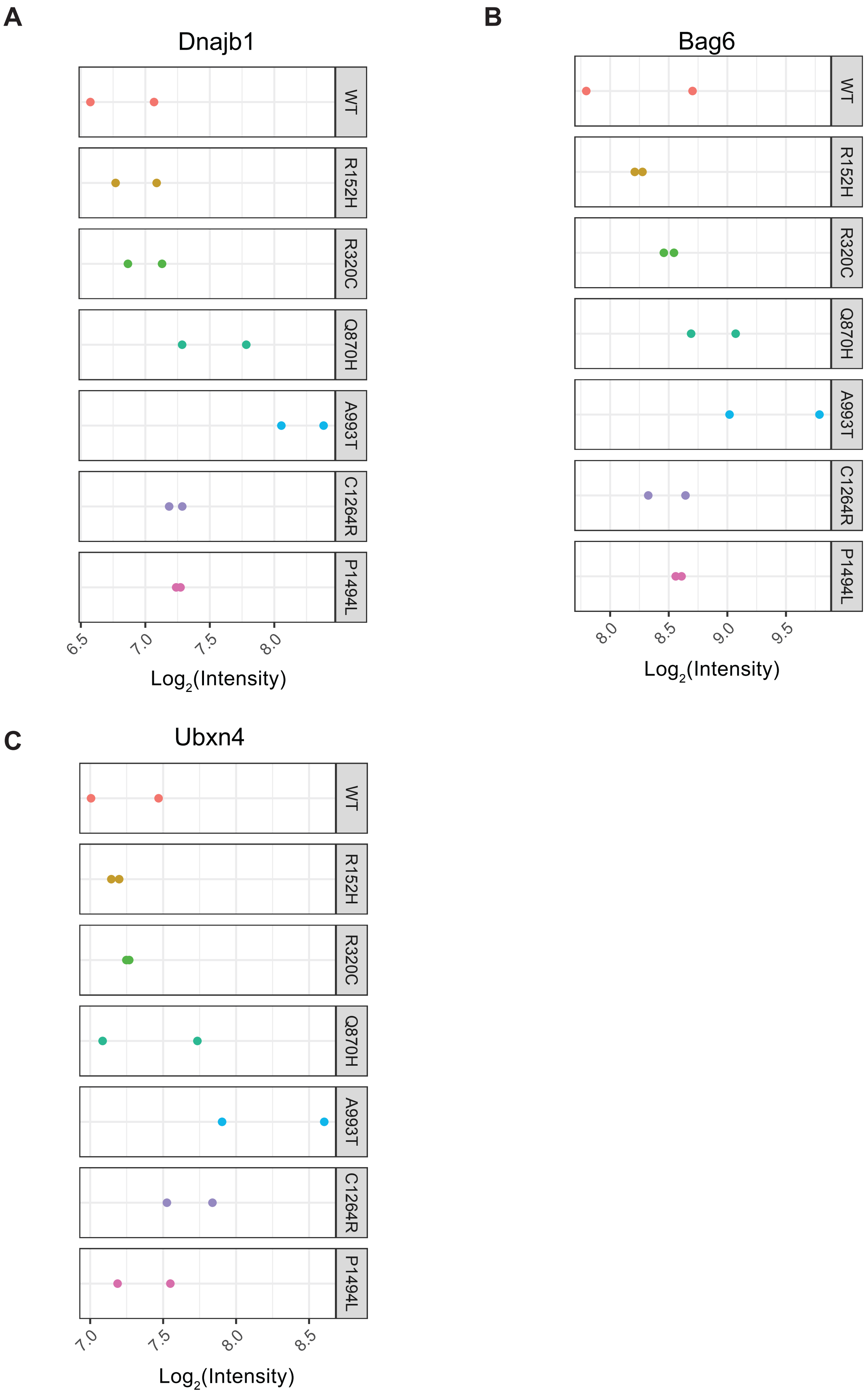


**Figure S7.** $Log_2$ intensities of **A)** Dnajb1, **B)** Bag6, **C)** Ubxn4 across all MS runs and variants.
